# Temporal resolution of melanogenesis determine fatty acid metabolism as key skin pigment regulator

**DOI:** 10.1101/2021.04.06.438588

**Authors:** Farina Sultan, Reelina Basu, Divya Murthy, Manisha Kochar, Kuldeep S. Attri, Ayush Aggrawal, Pooja Kumari, Pooja Dnyane, Archana Singh, Chetan Gadgil, Neel S. Bhavesh, Pankaj K. Singh, Vivek T. Natarajan, Rajesh S. Gokhale

**Affiliations:** CSIR-Institute of Genomics and Integrative Biology, New Delhi, 110025, India; Academy of Scientific and Innovative Research, Ghaziabad, UP, 201002, India; National Institute of Immunology, New Delhi, 110067, India; Eppley Institute for Research in Cancer and Allied Diseases, University of Nebraska Medical Center, Omaha, NE, 68198, United States; International Centre for Genetic Engineering and Biotechnology, New Delhi, 110067, India; CSIR-National Chemical Laboratory, Pashan, Pune, Maharashtra, 411008, India

## Abstract

Therapeutic methods to modulate skin pigmentation has important implications for skin cancer prevention and for treating meta-inflammatory-triggered cutaneous conditions. Modulators of cAMP signalling of melanocyte have met with minimal clinical efficacy. Towards defining new potential targets, we followed temporal dynamics orchestrating melanocyte differentiation by using a cell-autonomous pigmentation model. Our study elucidates three dominant phases of synchronized metabolic and transcriptional reprogramming. The induction phase is concomitant with a paradoxical decrease in MITF levels, reduced proliferation, and increased anabolic metabolism mediated by AKT phosphorylation. The melanogenic phase shows rapid uptake of glucose and fatty acid, transiently forming lipid droplets through SREBF1-mediated regulation of fatty acid metabolism. This heightened bioenergetic activity impairs mitochondria and the recovery phase is marked by a shift to aerobic glycolysis and activation of the NRF2 detoxication pathway. Finally, we show that inhibitors of lipid metabolism indeed resolve hyper-pigmentary conditions in a guinea pig UV-tanning model. Our studies reveal metabolic control mechanisms of melanocytes that could govern the balance between differentiation and proliferation in a variety of cutaneous diseases.

## Introduction

Despite significant progress in understanding the physiology and biochemistry of human skin pigmentation, the strategies to manipulate this phenomenon for clinical benefit has met with minimal success. To counter the deleterious effects of UV radiations, human skin activates melanisation that protects skin from cancer and photoaging ^1^. On the other hand, pathological hyperpigmentation response of skin occurs due to inflammatory conditions ^2^. Patchy cutaneous hyperpigmentation is associated comorbidity in more than 30% of patients with diabetes and obesity ^3^. Skin pigmentation is due to the presence of melanocyte cells in the epidermis, which possesses biosynthetic machinery to produce melanin within melanosomes ^4^. These specialized membrane-bound organelles are then transferred to neighbouring keratinocytes imparting photoprotection ^5, 6^. Melanogenesis thus can be considered to be a conglomeration of many interacting components wherein individual components as well as the interaction networks manifest spatiotemporal coherence. Perturbations within any of these events result in homeostatic imbalance leading to a disease phenotype. The challenge is to elucidate temporal dynamic interactions between constituent molecules of various cellular processes.

MITF, the central transcription regulator of melanocyte lineage, connect array of gene networks pertaining to melanogenesis, proliferation and survival ^7^. MITF regulates the expression of numerous pigmentation-associated genes such as PMEL17 and MART1 and melanin synthesis enzymes TYR, DCT and TYRP1 to promote melanocyte differentiation ^8–10^. Cellular homeostatic genes, including genes involved in apoptosis (eg, BCL2) and the cell cycle (eg, CDK2) are also regulated downstream to MITF ^11, 12^. UV-mediated activation of pigmentation proceeds through secretion of α-MSH by keratinocytes, which binds to epidermal melanocytes receptor, the melanocortin 1 receptor (MC1R), triggering cAMP production and CREB-mediated MITF transcription ^13–16^. Further, co-activators like SOX10 and Hippo signalling effector proteins activate MITF, while suppressors like TCF4 and ATF4 downregulate MITF transcriptional response ^17–19^. Melanocytes thus possess the ability to respond to environmental signals and assume a wide variety of distinct functional fates. Pharmacological inhibitors employed to modulate pigmentation are mostly designed to target cAMP pathway ^20^. However, taking them from cell culture studies to the clinical setting has been a challenging task. This could be due to the interaction of multiple signalling pathways that govern stages of melanogenesis including melanosome biogenesis, melanin synthesis and their transfer to neighbouring keratinocytes. These specialized cells of epidermis return to a resting state, where they are known to persist, potentially prepared for another round of activation. These distinct phases of melanocytes can be anticipated to be dependent on dynamic changes in cellular metabolism. Understanding how metabolism influence melanogenesis will open new avenues to identify targets for modulating pigmentation.

Several lines of evidence suggest the role of mitochondria in the induction of melanin synthesis, some of which are rather paradoxical. For example, induction of melanin synthesis in B16F10 melanoma cells reduces the oxygen consumption after 48 h of stimulation, without changes of mitochondrial membrane potential ^21^. However, mitochondrial mass is reported to be higher in cells with melanogenesis stimulation ^21^. A fraction of these mitochondria is shown to be in direct contact with melanosomes, where the inter-organelle connections mediated by fibrillar bridges are proposed to facilitate the exchange of small molecules between the two organelles ^22^. Remodelling mitochondria towards increased fission enhances ROS levels that can shut down melanogenesis ^23^. Contrary to this, targeting F1F0-ATP synthase that should also result in ROS accumulation, induces hyper-pigmentation ^24^. In a recent study, untargeted metabolomics of α-MSH induced B16F10 cells analysed at 1 hour, 24 hours and 48 hours showed minimal changes in the metabolite profile, when compared with their respective controls ^25^. While surprising, it is possible that the time points used in this study do not capture the dynamics of metabolite concentrations and metabolic fluxes. Another distinct possibility is the heterogeneity within cellular populations and cell surface receptors, which could obscure the interpretation of the results. Cellular heterogeneity has been reported for primary melanocyte cultures, where the cells could be transiting between precursor cells and their descendants at different stages of differentiation. For identification of transient regulatory events during melanogenesis, it is pertinent to resolve melanocyte function over time in a synchronized model system that can capture a full array of events.

With metabolic reprogramming is reemerging as a hallmark of cellular effector function, it is important to leverage metabolic dependencies as a possible target for modulating skin pigmentation. In this study, we have employed a previously developed B16 cell-autonomous pigmentation model where cells transit from basal depigmented to the pigmented state over a period of 6 days ^26^. Transcriptomic and metabolomic studies, allow us to identify dynamically changing key transcriptional network modules and corresponding metabolic configuration in a time-dependent manner. Along with defining a framework for understanding melanogenesis programming, our studies identify the transient, yet key role of SREBF1-mediated fatty acid metabolism during the melanogenic phase. Based on the guinea pig tanning model, we show that inhibitors of fatty acid metabolism can resolve hyper-pigmentary conditions, thus revealing new targets for modulating skin pigmentation.

## RESULTS

### Melanocyte differentiation is coupled to transcriptional regulation of metabolic networks

To understand the differentiation programming of melanocytes from depigmented to pigmented state, we perform global transcriptomic analysis of B16 cell-autonomous pigmentation model ^26^. In this model, a transition from basal depigmented to the pigmented state of cells occurs over a period of 6 days and is triggered by the fine balance between the intrinsic needs of the cells and the constraints imposed by the extrinsic conditions. As phenotypic changes in the melanin are best observed and quantitated from day 3 to day 6 (Sup Fig 1A), these time points are taken for temporal analysis of melanocyte differentiation. Principal component analysis (PCA) of the transcriptome data displayed tight clustering for different biological replicates indicating relatedness while segregating the day-wise clusters (Sup Fig 1B). We then performed differential expression analysis using DESeq2, with an adjusted p-value < 0.001 (Fig 1A). Amongst the differentially expressed genes, several key pigmentation-related genes like *Mitf*, *Pmel, Tyr* could be observed. Previous studies have suggested a decrease in MITF levels leading to the activation of pigmentation. Indeed, we observe an analogous trend of drop in *Mitf* expression, while the downstream target genes of *Mitf,* i.e., *Pmel, Tyr* levels increase from day 3 to day 6, (Sup fig 1C, 1D). Further, we observe a linear decrease in proliferation rate during these four successive days, with about 40% decrease observed on day 6, as compared to day 3 (Sup Fig 1E). As expected, the cell cycle gene CDK2 also shows significant downregulation (Sup Fig 1F, 1G). Hierarchical clustering analysis of 1493 genes, identified based on p-value < 0.001, were segregated into seven clusters. In general, genes on day 3 and day 4 show substantially similar gene expression values as compared to day 5 and day 6. We then performed pathway enrichment analysis for these gene clusters by using the KEGG database, illustrated as Bubble Plot (Fig 1B). Pathways like RNA transport, ribosome biogenesis, and spliceosome were enriched on day 3 and day 4, indicative of transcriptional activation during early pigmentation phases. Day 5 showed upregulation of metabolic pathways like steroid and unsaturated fatty acid synthesis and fatty acid metabolism, while day 6 displays an up-regulation of glycolysis and glutathione metabolism. The strikingly differential enrichment observed for metabolic pathways during different days suggested that dynamic changes in metabolism could influence pigmentation.

**Figure 1:**
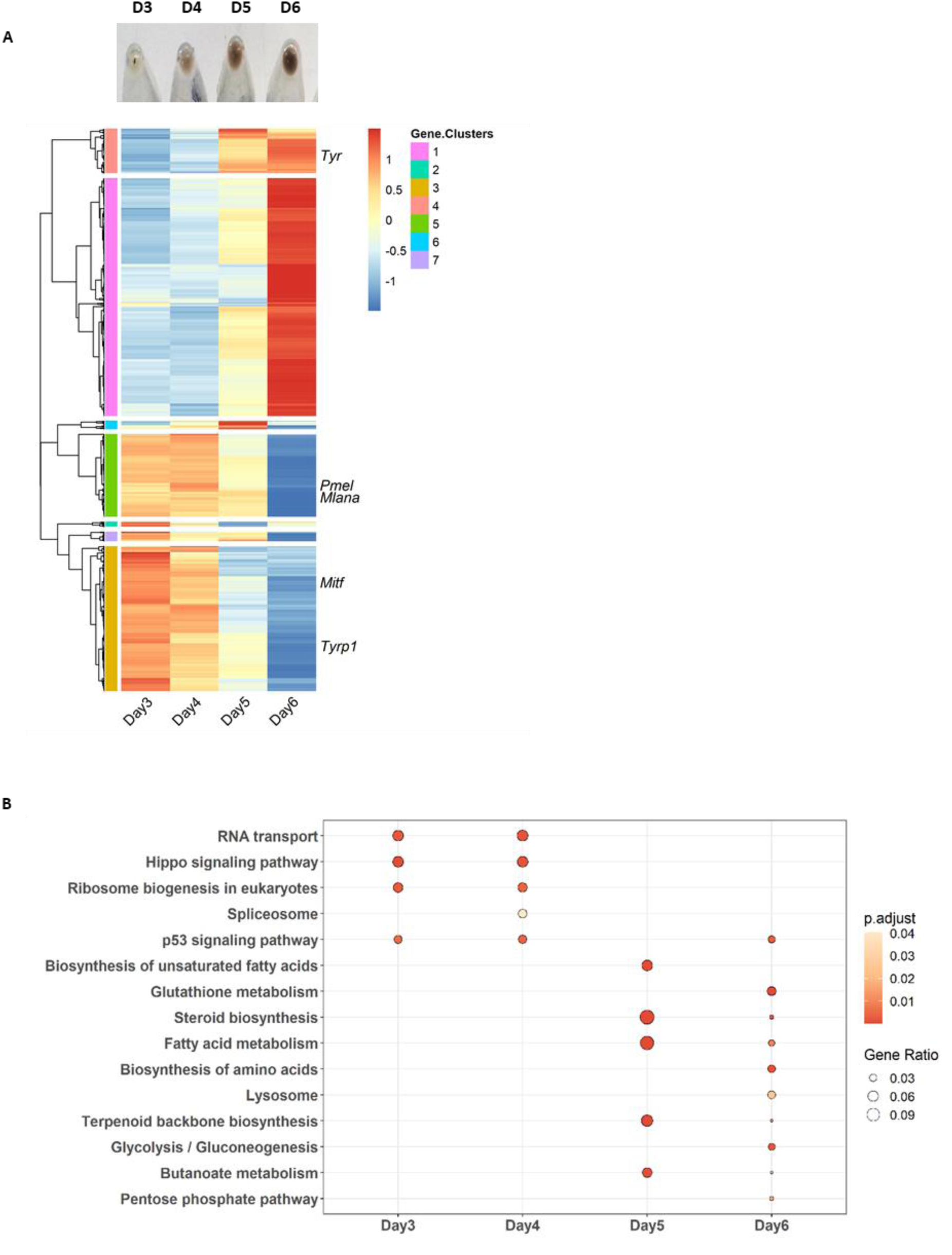
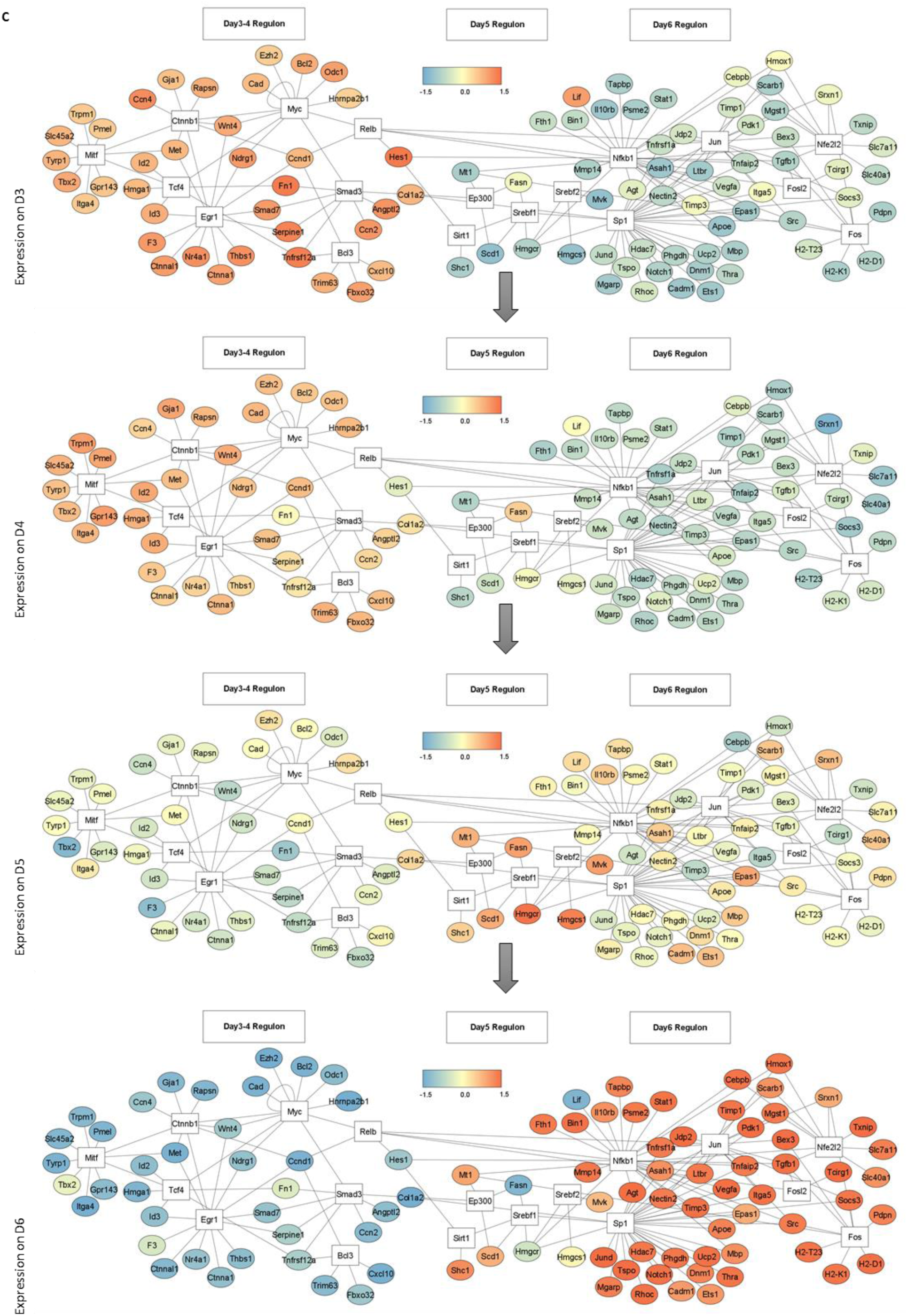
**Melanogenesis is coupled to transcriptional regulation of metabolic networks** A. Heatmap represents the differentially regulated gene from day 3 to day 6 obtained in Deseq2 analysis with adjusted p-value < 0.001, for two independent biological replicates. Scale from blue to red represents z-score for fold change from -1 to +1. Hierarchical clustering was done based on expression similarity. B. KEGG pathway enrichment analysis was done for upregulated genes on day 3, 4, 5 and 6 obtained by Deseq2. Bubble plot depicts the enrichment of pathways on different days, where the size of bubble represents the gene ratio and colour represents the p- value. C. TF-TG network analysis for top 7 upregulated transcription factors along with the set of target genes on day 3, 4, 5 and 6. TF are enclosed in white box and the line connect each TF to the set of genes regulated by them during pigmentation. Target genes are enclosed in circle and expression is represented by the scale from red to blue.

To identify prominent transcriptional networks on day 3, day 4, day 5 and day 6, we performed transcription factor enrichment analysis for differentially regulated genes using the TRRUST database. We constructed transcription factor (TF)-target gene (TG) network maps for the top 7 transcription factors identified on each day. We then overlaid the changes in transcript levels of target genes to generate temporal expression plots (Fig 1C). This representation elucidated an interesting representation of the different TF-TG network maps during the course of pigmentation. On day 3, pigmentation regulators *Mitf* and *Egr1* networks are functional, both of which are known to induce pigmentation response in melanocytes ^27, 28^. Another melanogenesis-associated gene *Tcf4* could also be noted in the day 3-4 regulons, which is known to suppress *Mitf* levels ^18^. It is interesting to note that *Tcf4* TG expression increases on day 4 as compared to day 3, suggestive of a possible role in dampening of MITF levels. The expression of *Myc* network, which is involved in proliferation, decreases with each day and this is congruent with our experimental observations ^29^. On day 5, completely new sets of TF-TG networks emerge, prominent of them are the *Srebf1* and *Srebf2* clusters, which are known to regulate lipid metabolism across different cell types ^30^. We examined mRNA expression of *Srebf1* and its key downstream targets like *Fasn*, *Acaca*, *Acacb,* and *Acly* by qRT-PCR analysis. About 1.5- to 2-fold changes could be noted for all four TGs on day 5 and day 6 (Sup Fig 1H, 1I), corroborating our computational analyses. FASN upregulation was also confirmed at the protein level (Sup Fig 1J, 1K). Another significant cluster that becomes evident is the *Nfe2l2 (Nrf2)* regulon. This transcription factor is involved in the phase II detoxification pathway and previous studies have shown that melanocytes resist oxidative detoxification through a robust expression of this pathway during pigmentation ^31^. Together, the transcriptional analysis provides interesting insights into the temporal regulation of TF-TG networks and pathway analysis suggest a modulatory role for metabolism during these different phases of melanocyte differentiation.

### Steady-state analysis of polar metabolites using mass spectrometry during pigmentation

To understand how metabolic changes are linked with the process of melanogenesis, we measured the levels of polar metabolites from day 3 to day 6 cells using liquid chromatography-coupled tandem mass spectrometry. A total of 306 peaks were obtained, out of which 175 were mapped to different metabolites with high confidence. Relatedness between all the data sets was compared using PCA. All replicates showed good clustering and clear day-wise segregation was observed (Sup fig 2A). Hierarchical clustering analysis based on the top 50 regulated metabolites showed an increased level of metabolites corresponding to nucleotide and amino acid metabolism during day 3 and day 4. On day 5 and day 6, a substantial number of cofactors and Kreb’s cycle metabolites were seen to be up-regulated (Sup Fig 2B). To understand the pathway-based connectivity, we overlaid the data on four major central carbon metabolic pathways- glycolysis, tricarboxylic acid cycle (TCA), pentose phosphate pathway (PPP), and hexosamine biosynthesis pathway (HBP) (Fig 2A). Analysis of glycolytic intermediates showed two distinct patterns of regulation. Metabolites in the upper half of the glycolysis (six-carbon metabolites) showed limited variations across all four days. The upper half of glycolysis is known to branch into the PPP and HBP. Indeed, analysis of metabolites from PPP and HBP indicated an increased accumulation on day 5 and day 6. PPP generates NADPH to maintain a redox environment and provides intermediates for fatty acid and nucleotide synthesis. HBP forms uridine diphosphate-β-N-acetylglucosamine (UDP-GlcNAc) moiety required for O-linked β-N-acetylglucosamine (O-GlcNAc) posttranslational modification of melanogenic proteins ^32^. The metabolites in the lower half of glycolysis (three-carbon metabolites) increased ∼4-fold during the pigmentation phase. The lower half of the glycolysis produce lactate and feeds into the TCA cycle. Interestingly, the levels of both lactate and TCA metabolites show an increase with time. Together, these steady-state metabolic analysis suggests that the glucose is effectively distributed into all the branches of central carbon metabolism during pigmentation. It is likely that this alteration in the metabolic pools supports the effector functions of melanocytes. It is, therefore, apparent that melanogenesis activation is not accompanied merely by a switch from oxidative metabolism to glycolysis-TCA, but that both pathways are upregulated to support bioenergetic demands. Furthermore, increased HBP and PPP pathway metabolism suggests support for the synthesis of a variety of biomolecules.

**Figure 2:**
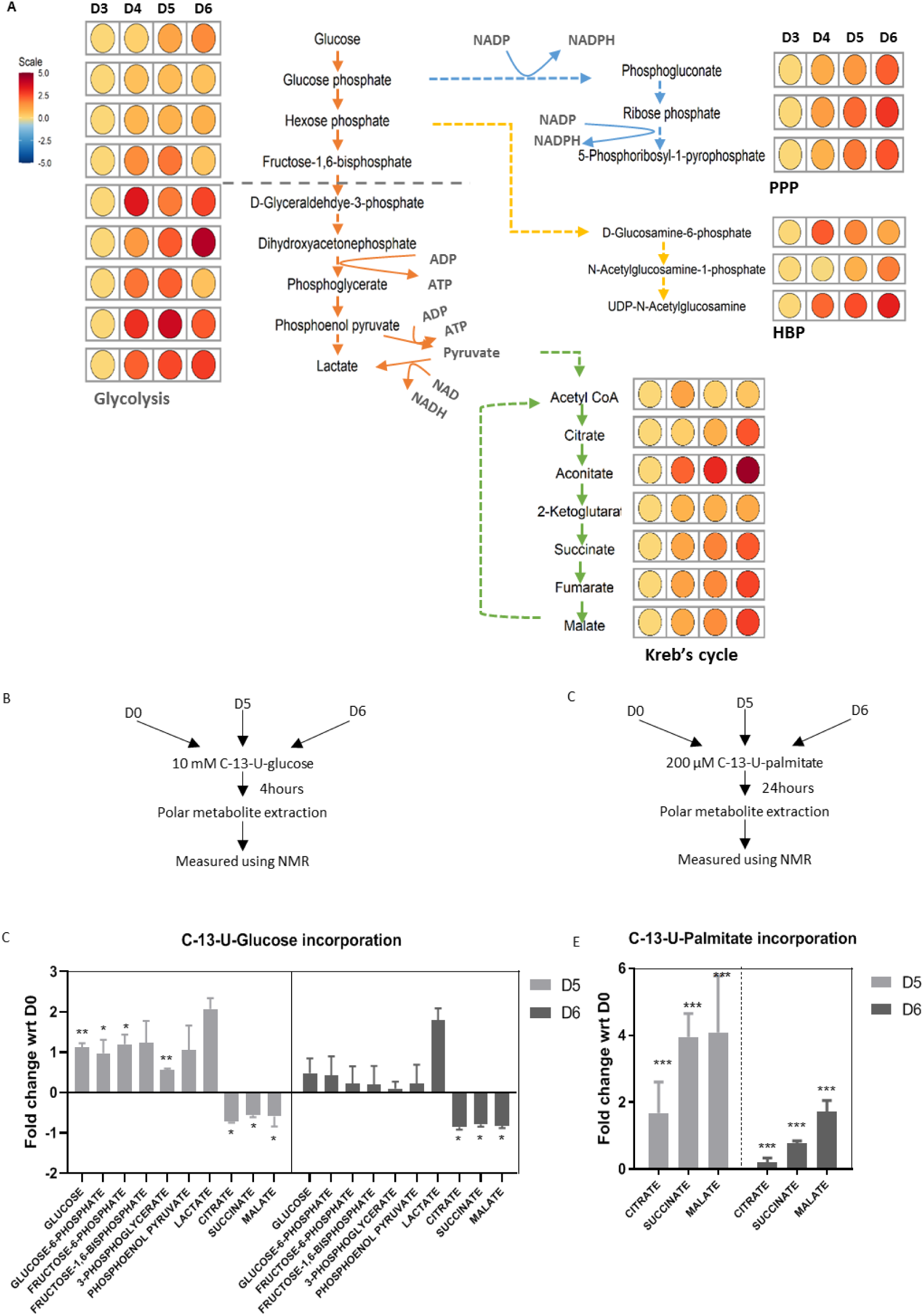
**Partial uncoupling of glycolysis and TCA supports biosynthesis pathways (PPP, HBP) while FAO increases to sustain TCA** A. Liquid chromatography-coupled tandem mass spectrometry-based metabolomic analysis is performed for polar metabolites extracted from D3 to D6 for four independent replicates. Metabolic network is drawn for Glycolysis, Kreb’s cycle, Pentose phosphate pathway (PPP) and Hexosamine biosynthesis pathway (HBP) by comparing their metabolite levels across D3 to D6. Fold changes are plotted as heatmaps along each metabolite in the pathway with respect to D3. Scale bar ranges from blue to red corresponding to change from -5 to +5 fold. B. Schematic depicting study design for pulse-chase labelling of uniformly ^13^C labelled glucose in glycolysis and TCA metabolites at D0, D5 and D6 using NMR. C. NMR-based quantitation of [U-^13^C]-glucose incorporation in Glycolysis and TCA metabolites at D5 and D6 with respect to D0. Mean ± s.e.m. is plotted for three independent replicates. Two-way ANOVA is applied F (1.667,33.33) = 33.95, p-value < 0.0001. Dunnett’s test is performed for pairwise analysis. ***p-value < 0.005, **p- value < 0.01, *p-value < 0.05. ns is not significant. D. Schematic depicting study design for pulse-chase labelling of uniformly ^13^C labelled palmitate in TCA metabolites at D0, D5 and D6 using NMR. E. NMR-based quantitation of [U-^13^C]-palmitate incorporation in TCA metabolites at D5 and D6 with respect to D0. Mean ± s.e.m. is plotted for three independent replicates. Two-way ANOVA is applied F(2,12) = 19.91, p-value < 0.0002. Dunnett’s test is performed for pairwise analysis. ***p-value < 0.005, **p-value < 0.01, *p-value < 0.05. ns is not significant.

### [U-^13^C]-Palmitate incorporation in TCA metabolites during pigmentation phase

To understand how the glucose uptake during the pigmentation phase (day 5 and day 6) is quantitively apportioned between the lactate and TCA metabolites, we performed stable isotope tracing using labelled [U-^13^C]-Glucose. These pulse-labelling experiments were analyzed after 4 hours of addition of 10 mM [U-^13^C]-Glucose in the glucose-free medium (Fig 2B). The incorporation of ^13^C isotope in different metabolites was followed by NMR-based measurements and several metabolites could be identified with high confidence. We observed a two-fold increase in the ^13^C labelling of glycolytic metabolites on day 5 when compared with depigmented cells (Fig 2C). In contrast, the TCA metabolites- citrate, succinate, and malate - showed a substantial decrease in the ^13^C incorporation. This configuration of increased glycolytic metabolites and decrease in TCA metabolites is also observed for day 6, suggesting that the pigmentation phase is associated with decreased channelization of pyruvate to acetyl CoA. Interestingly, incorporation of ^13^C labelling on day 6 in glycolytic metabolites is lower than day 5, while relatively little variation is observed for TCA metabolites and lactate levels. This suggested an increased uncoupling between glycolysis and TCA. Enhanced glycolysis flux on day 5 and day 6 is majorly contributed by increased utilization of glucose from media (Sup Fig 2C). Pyruvate dehydrogenase kinase (PDK1) can regulate the activity of pyruvate dehydrogenase (PDH), which converts pyruvate to acetyl-CoA ^33, 34^. We indeed observed increased expression of *Pdk1* during the late phase of pigmentation that substantiates the quantitative increase in the level of lactate observed during day 6 (Sup Fig 2D, 2E, 2F). Overall, this indicates uncoupling of glycolysis and TCA in the late recovery phase of pigmentation. However, we observed an increased accumulation of TCA metabolites in the steady-state analysis, which suggested the possibility of significant carbon flux is being channelized from alternate pathways to TCA. Several studies have shown that cells can compensate for decreased glucose flux by utilizing glutamine and fatty acids to maintain the TCA cycle functions ^35, 36^. In several processes, fatty acid oxidation has emerged as an alternative source of generating acetyl-CoA pools, particularly during effector functions ^37, 38^. Incidentally, B16F10 cells cannot grow in the absence of glutamine and we, therefore, scrutinized whether [U-^13^C]-Palmitate could be channelled into TCA metabolites during the pigmentation phase (Fig 2D). NMR measurements of ^13^C label incorporation revealed a more than 2-fold increase in the incorporation on day 5 when compared with depigmented cells (Fig 2E). This incorporation again decreases on day 6, suggesting a diminishing role of oxidative phosphorylation in maintaining the energy demands of the cells during the late phase of pigmentation. Such a dynamical metabolic adaptation during the pigmentation phase may be a crucial metabolic switch that shunts fatty acids to mitochondria to maintain ATP production and sustain PPP via glycolytic shunting to keep a balance between available nutrients and energy balance.

### Fatty acid availability increases mitochondrial respiration during pigmentation

To determine how cellular bioenergetics is affected by the availability of carbon source, we measured cellular oxygen consumption rate in either glucose- or oleate-supplemented media using the Seahorse Mito-stress assay (Fig 3A). We observed that the basal respiration rate, which accounts for total oxygen consumption by the cells, is higher on day 5 than on day 6 (Fig 3B). However, in the presence of oleate, cells show higher basal respiration on day 5 in comparison to glucose. Availability of oleate in the media also results in higher spare respiration capacity of cells on both days, day 5 and day 6 (Fig 3C). This indicates that increased electron flow through mitochondrial complexes reaches a maximum on day 5 when fatty acids rapidly undergo β-oxidation to provide more reducing equivalents for the electron flow in the electron transport chain (ETC) than glucose. Together, these studies provide support to the hypothesis that glucose is partially uncoupled from TCA while fatty acid oxidation is the preferred pathway during the intermediate phase of pigmentation. The decrease observed on day 6 for mitochondrial respiration may be due to altered mitochondrial structure, which is essential to maintain respiratory complex protein in close proximity for transporting electrons in ETC ^39^. To test this possibility, we performed transmission electron microscopy (TEM) based ultrastructural analysis of mitochondria during melanogenesis (Fig 3D). Our data showed the presence of a few stage III/IV (electron-dense) melanosomes on day 3 and day 4, which dramatically increases up to 6-8 times on day 5 and day 6, respectively (Fig 3E). At the same time, we also observed an increase in the number of unhealthy mitochondria during the late pigmentation phase, as determined by cristae stacking and the presence of vacuoles around mitochondria. The number of unhealthy mitochondria observed is about 40% on day 6, which could substantially impact mitochondrial respiration (Fig 3F). It is clear that during pigmentary cascade, fatty acid metabolism is crucial to support energy metabolism, while in hyper- pigmented cells, cumulative stress due to high OXPHOS and melanogenesis, with enhanced ROS, results in an increased percentage of damaged mitochondria.

**Figure 3:**
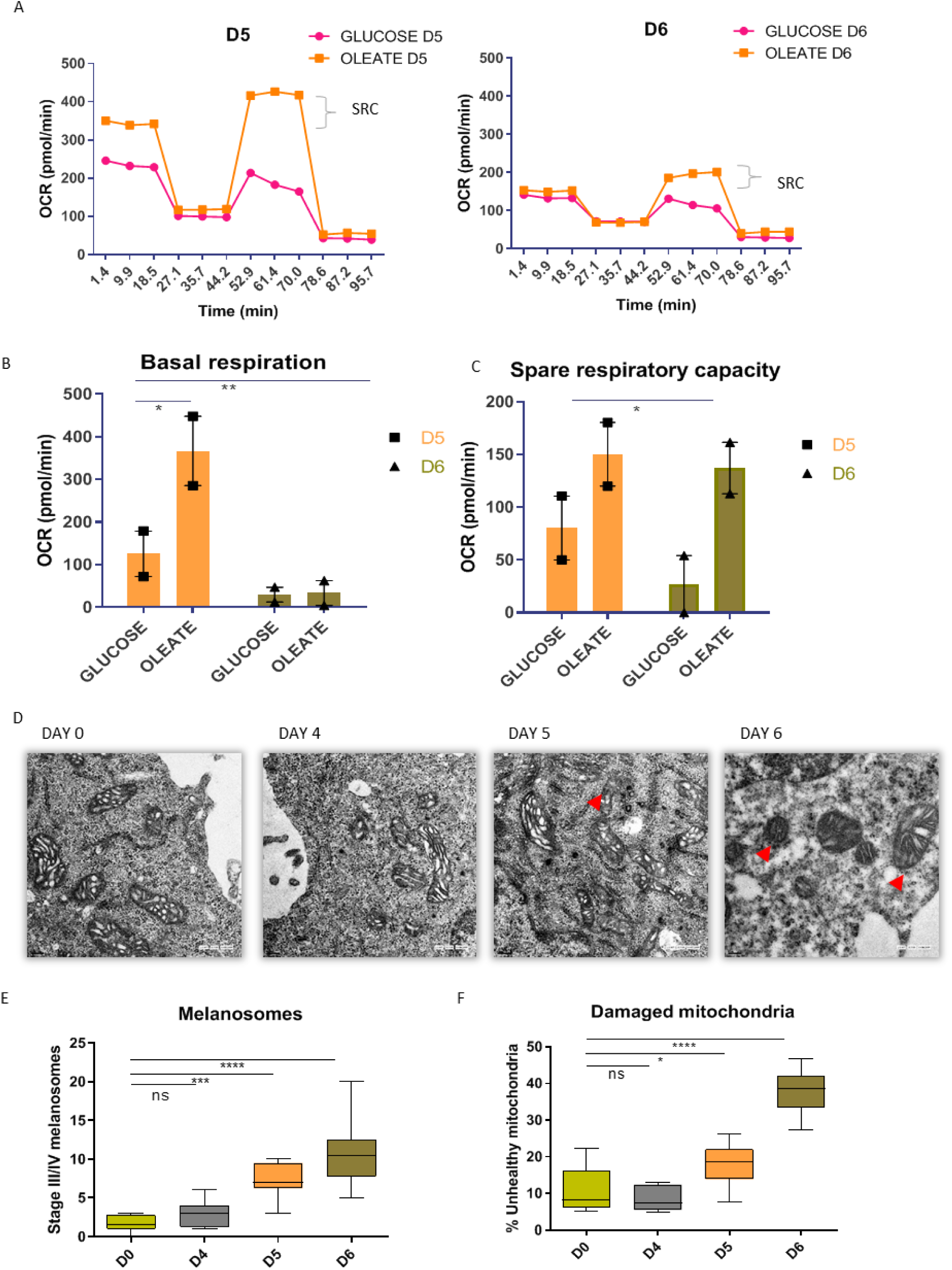
**Fatty acids are the preferred carbon source for mitochondrial respiration during pigmentation** A. Comparative analysis of oxygen consumption rate in glucose or fatty acid supplemented media on D5 and D6 using Seahorse Mito stress assay. Represented plot depicts median value for two independent biological replicates with three technical replicates in each set. B. Quantitative analysis of basal respiration in glucose or fatty acid supplemented media on D5 and D6. Mean and Range is plotted for two biological replicates with three technical replicates for each set. One-way ANOVA is applied F (5,8) = 9.693, p-value = 0.003. Tukey’s test is performed for pairwise analysis, **p-value < 0.01, *p-value < 0.05. C. Quantitative analysis of spare respiratory capacity in glucose or fatty acid supplemented media on D5 and D6. Mean and Range is plotted for two biological replicates with three technical replicates for each set. One-way ANOVA is applied F (5,9) = 3.792, p-value = 0.0399. Tukey’s test is performed for pairwise analysis. **p- value < 0.01, *p-value <0.05. D. Transmission electron microscopy-based analysis showing mitochondrial cristae morphology change during pigmentation. Images were taken at 3500X magnification. E. Box plot depicting quantitative analysis of Stage III/IV melanosomes on different days (Number of cells, n = 8). Whiskers represent min and max range with mean as center line. One-way ANOVA is applied F (3,28) = 19.60, p-value < 0.0001. Tukey’s test is performed for pair-wise comparison. ****p-value < 0.0001, ***p-value =0.007. ns is not significant. F. Box plot depicting quantitative analysis of damaged mitochondria on different days (Number of cells, n = 8). Whiskers represent min and max range with mean as center line. One-way ANOVA is applied F (3,28) = 47.08, p-value < 0.0001. Tukey’s test is performed for pair-wise comparison. ****p-value < 0.0001, *p-value = 0.027. ns is not significant.

Altogether, from transcriptomic and metabolomics analysis, we propose that melanogenesis is divided into three phases – preparatory, melanogenic and recovery phase. Day 3 and day 4 captures the preparatory phase where the MITF mediated signalling networks are induced and anabolic pathways are activated. Followed by the melanogenic phase on day 5 where pigmentation synthesis machinery is high and metabolic changes like increased fatty acid utilization occurs to support pigmentation. Day 6 profile captures recovery pathways where pigment inhibitory pathways and recovery mechanism are upregulated.

### Fatty acid metabolism is the critical pathway in mediating pigment production

After recognizing that cells can potentially utilize fatty acids to sustain ATP production during the pigmentation phase, we were interested to understand the mechanisms through which these cells rewire fatty acid metabolism. As free fatty acids are toxic to cells, an important facet of lipid metabolism is the assimilation of neutral lipids (triacylglycerols) as lipid droplets. We traced neutral lipid content in B16 cells during pigmentation by using BODIPY dye (Fig 4A). Quantitative analysis of number as well as the volume of lipid droplets revealed the formation of these lipid aggregates increases early on D5 (Fig 4B, 4C). Further, these lipid droplets were rapidly depleted on D6. These results suggest that B16 cells assimilate fatty acids synthesized by cells as lipid droplets and probably utilized for energy generation during pigmentation.

**Figure 4:**
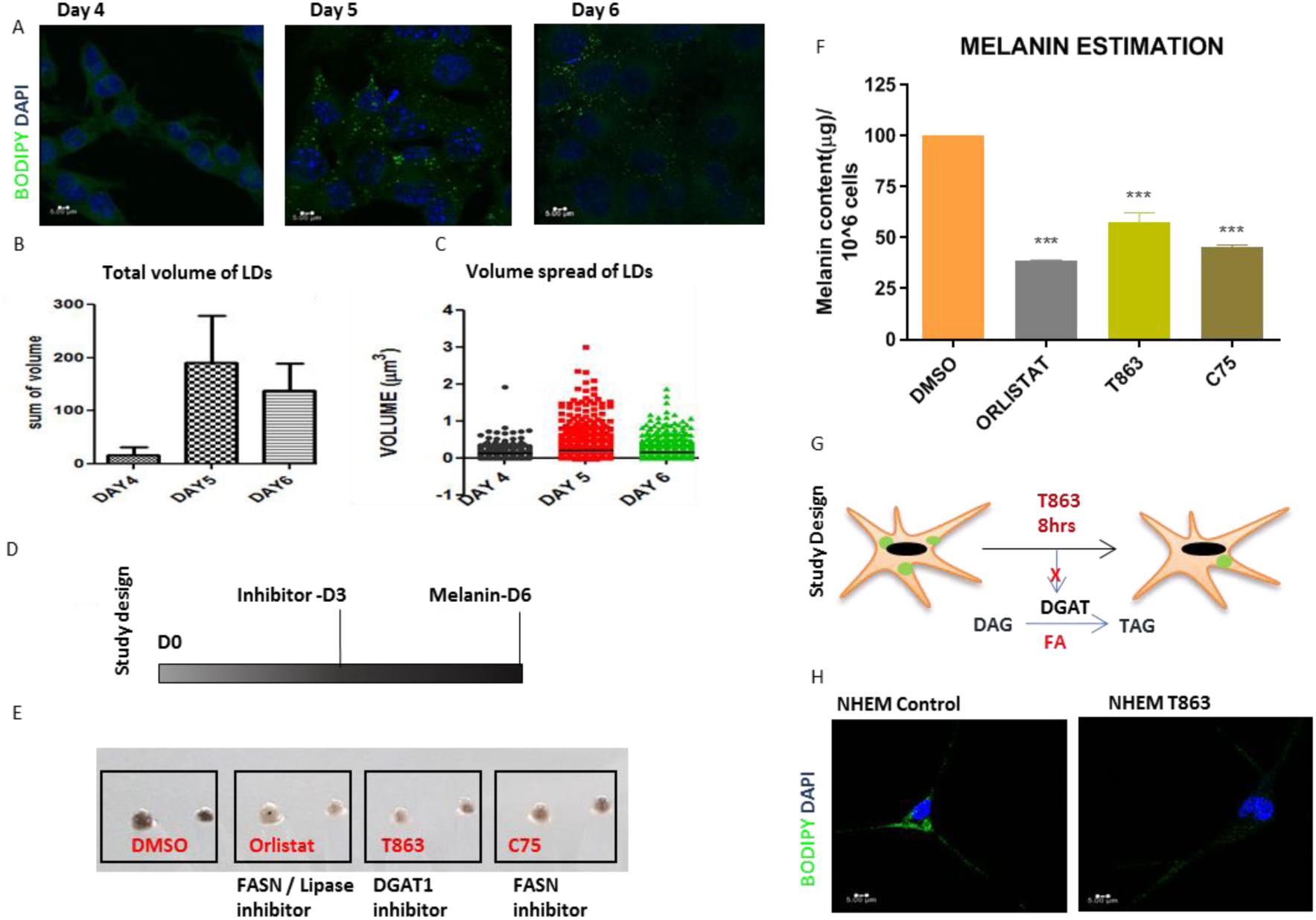
**Fatty acid metabolism is a critical pathway in mediating pigment production** A. Representative confocal images showing lipid droplet accumulation in B16 cells during pigmentation. Images were taken at 60X. Scale is 5µm. B. Bar graph depicting the quantitation of total volume of lipid droplets using VOLOCITY software. Approximately 50 cells are taken in three independent biological replicates. C. Dot plot depicting the quantitation of size of lipid droplets using VOLOCITY software. Approximately 50 cells are taken in three independent biological replicates. D. Schematic showing experimental timeline for inhibitor addition and analysis of pigmentation in B16 cells. E. Representative image showing phenotypic differences in melanin accumulation upon inhibitors treatment in two technical replicates of D6 cells. Two technical replicates are taken for three independent biological sets. F. Bar graph depicting melanin estimation after different inhibitor treatment. Mean ± s.e.m. is plotted for three biological replicates. One-way ANOVA is applied F (3,4) = 149.9. p-value = 0.0001. Dunnett’s test for pair-wise comparison. ***p-value < 0.005, **p-value < 0.01, *p-value < 0.05. G. Schematic study design for analyzing triacylglycerol formation in primary melanocyte culture by inhibiting DGAT1 using T863, and capturing lipid droplets using BODIPY dye. H. Representative images showing lipid droplet accumulation in primary melanocytes upon T863 addition. Images were taken at 60X. Scale is 5µm.

To ask whether fatty acid metabolism is important for pigmentation, we performed an inhibitor screen for specific enzymes involved in *de nov*o fatty acid synthesis, TAG synthesis and lipolysis. B16 cells were treated with FASN inhibitor (C75-20 µM), DGAT inhibitor (T863- 10 µM), and lipase inhibitor/ FASN inhibitor (Orlistat-50 µM) on day 3 and analyzed for pigmentation differences on day 6 (Fig 4D). As expected, C75 and T863 decrease lipid droplet content of the cells on day 5, while Orlistat results in increased accumulation of lipid droplets suggesting that it acts as a lipase inhibitor (Sup Fig 3A). We noted a substantial decrease in the pigmentation for all three inhibitors, with 60% on Orlistat treatment, 40% on T863 treatment and 50% on C75 treatment (Fig 4E & F). The melanin estimation data manifest at the molecular level, as measured by tyrosinase protein levels (Sup Fig 3B, 3C). These studies reemphasize the significance of fatty acid metabolism during pigmentation.

We validated these findings of lipid droplet formation in the pigmented primary melanocytes. BODIPY staining showed abundant lipid droplets in cultured primary melanocytes that increased noticeably on oleate feeding (Sup Fig 3D). Further, we probed whether T863, an inhibitor of DGAT1 that catalyzes the rate-limiting step of diacylglycerol (DAG) to triacylglycerol (TAG) formation, could potentially alter the levels of lipid droplets (Fig 4G). Treatment with T863 for 8 hours resulted in the complete disappearance of the lipid droplets indicating a high turnover of lipids in pigmented melanocytes (Fig 4H).

While primary melanocytes isolated from neonatal foreskin sample from Indian origin are highly pigmented and have abundant lipid droplets, B16 cells are depigmented initially and transiently accumulate lipid droplets during the transition from depigmented to pigmented state. The above studies highlight the importance of *de novo* lipogenesis and the storage of lipid droplets as an integral feature of melanogenesis. The exogenous uptake of fatty acids by melanocytes, which is higher during the early and intermediate phase probably imparts metabolic flexibility to these cells for effective pigmentation (Sup Fig 3E).

### Epidermal melanogenesis mediator α-MSH activates SREBF1

As fatty acid metabolism is important during pigmentation and the SREBF1 network was amongst the prominently regulated network in TF-TG analysis, we interrogated the role of SREBF1 during melanogenesis. Towards this, we studied the effect of siRNA-based downregulation of *Srebf1* on pigmentation genes (Fig 5A). Transfections were carried out with Smart pool *Srebf1* siRNA on day 3 and transcriptional changes in pigmentation genes were monitored on day 5. We noticed about a 40-50% decrease in the *Srebf1* levels and similar downregulation was observed in its downstream target *Fasn* levels. Further, a significant decrease in expression of *Tyrp1* was seen, a gene involved in melanin synthesis, upon *Srebf1* downregulation. Next, we utilized 25-hydroxycholesterol (25-HC), a pharmacological inhibitor of SREBF1 activation, and evaluated its effect on pigmentation. The inhibitor was added on day 3, and analysis was performed on day 6. Along with the phenotypic change in pigmentation (Fig 5B), quantitative analysis showed a 50% decrease in tyrosinase expression upon 25-HC treatment (Fig 5C, 5D). These studies propose activation of SREBF1 during pigmentation.

**Figure 5:**
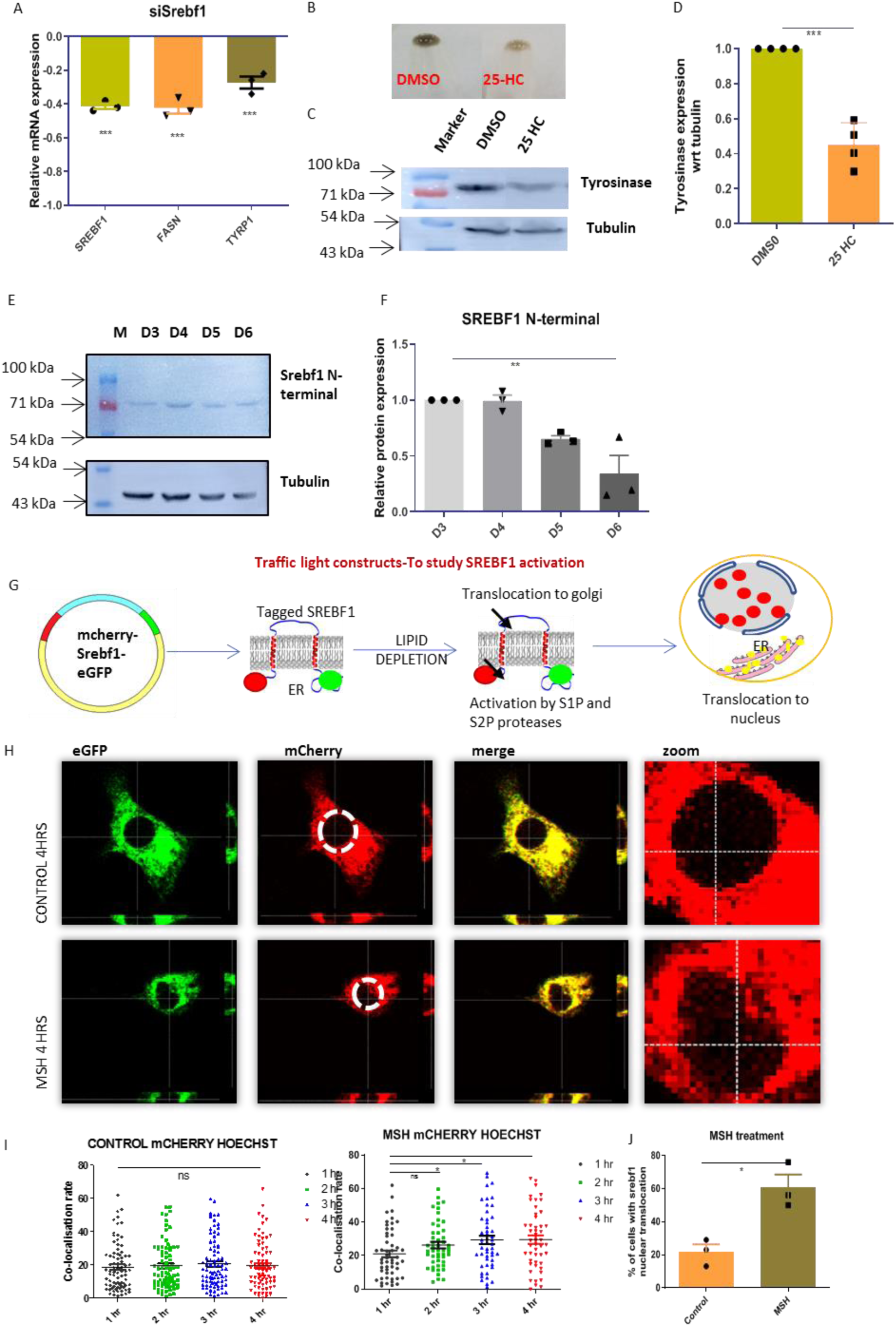
**Melanogenesis induction causes early activation of Srebf1** A. Bar graph representing qRT-PCR based quantitation of *Srebf1, Fasn* and *Tyrp1* genes on D5 upon silencing of *Srebf1* using smart pool siRNA. Mean ± s.e.m. is plotted for three biological replicates One-way ANOVA is applied, F (3,8) = 61.03, p-value < 0.0001. Tukey’s test is performed for pairwise analysis. ***p-value < 0.005, **p-value < 0.01, *p-value < 0.05. B. Representative B16 cell pellet images showing the phenotypic difference in melanin accumulation on day 6 upon inhibiting SREBF1 activation with 25-HC treatment. C. Representative western blot showing TYR protein expression, normalized to tubulin, upon 25-HC treatment. D. Bar graph depicting quantitative fold change of TYR expression with respect to control set. Mean ± s.e.m. is plotted for three independent biological replicates. Two-tailed unpaired t-test is performed, t=8.663, df=6. ***p-value =0.0001. E. Representative western blot showing SREBF1 N-terminal levels on different days of pigmentation, normalized to tubulin. F. Bar graph depicting quantitative fold change of Srebf1 N-terminal expression with respect to D3. Mean ± s.e.m. is plotted for three biological replicates. One-way ANOVA is applied, F (3,8) = 12.90, p-value = 0.002. G. Schematic design for studying SREBF1 activation using traffic light mCherry-Srebf1- eGFP vector and analyzing localization in different cellular compartments. Full-length protein localized to ER gives yellow fluorescence due to co-localization of mCherry and eGFP. Cleaved N-terminal is transported to the nucleus, thus, accumulation of mCherry in the nucleus is indicative of activation. H. Representative fluorescence images showing eGFP, mCherry and merge signal. Magnified nuclear images were shown to focus on SREBF1 nuclear translocation after 4 hours of MSH treatment. DMSO is taken as vehicle control. Around 30 cells were analyzed in each of the three biological replicates. One-way ANOVA is applied. For DMSO, F (3,325) = 0.2578, for MSH F (3,196) = 3.133, p-value < 0.0267. Tukey’s test is performed for pairwise analysis. ***p-value < 0.005, **p-value < 0.01, *p-value < 0.05. I. Dot plot depicts co-localization rate between mCherry and Hoechst signal analyzed for each cell at 1 hour, 2 hours, 3 hours and 4 hours, for control and MSH treatment. ns is not significant. J. Bar graph depicting the quantitation of number of cells showing positive phenotype after MSH treatment, determined by increased co-localization rate of mCherry and Hoechst signal from 1 to 4 hours. Student’s t-test is performed, t=4.267, df=4 p-value = 0.0130.

SREBF1 is an ER-resident protein that is activated upon the cleavage and nuclear translocation of its N-terminal domain. To examine SREBF1 activation, we performed western blot analysis with SREBF1 N-terminal antibody (Fig 5E). Quantitation of cleaved SREBF1 N-terminal band showed increased levels on day 4, which then slowly declined on day 5 and day 6 (Fig 5F). As SREBF1 activation is necessary for the induction of its downstream targets, we speculate that early activation of SREBF1 is important for efficient pigmentation. We then explored whether the classical inducer of melanogenesis, α-MSH, could activate SREBF1. To study this, we designed an activation assay for SREBF1 using a ‘traffic light construct’, wherein the mCherry tag was fused with the N-terminal of SREBF1 and eGFP was fused with C-terminal (Sup Fig 4). Upon SREBF1 activation, mCherry along with the N-terminal SREBF1 fragment would get translocated to the nucleus while the eGFP remains localized in ER (Fig 5G). Functional validation of this assay was assessed with insulin, which is a known activator of SREBF1 in hepatocytes ^40^ (Sup Fig 5A). Insulin indeed activated SREBF1 in melanocytes within hours in about 65% of the cells (Sup Fig 5B, 5C). Further, live-cell imaging was carried out for 4-6 hours after α-MSH treatment using confocal microscopy. Representative images show mCherry and eGFP signal in different channels after 4 hours of treatment (Fig 5H). Careful analysis for red signal in the nuclear region shows that the presence of mCherry increases after α-MSH treatment. Time-dependent changes in the co-localization rate of mCherry and Hoechst dye was analyzed after 1, 2, 3 and 4 hours to follow kinetic activation (Fig 6I). In the case of DMSO, the co-localization rate of mCherry and Hoechst dye did not increase in a time-dependent manner, suggesting no activation of SREBF1. For α-MSH, we observed that the co- localization rate of mCherry and Hoechst significantly increases after 3 hours. Further, the number of cells showing positive phenotype considerably increases to 60% after α-MSH treatment (Fig 6J). This data suggests that α-MSH induces nuclear translocation of SREBF1 N-terminal in B16 cells. Insulin is shown to activate SREBF1 via AKT pathway ^40^. In melanocytes, we observe that the phosphorylation of AKT is high on day 3 and decreases as days progress (Sup Fig 5D, 5E). We propose that SREBF1 activation in melanocytes could be mediated by AKT. This study proposes the role for the activation of SREBF1 on pigmentation induction, ensuring efficient fatty acid synthesis for heightened melanosome biogenesis and melanin synthesis.

**Figure 6:**
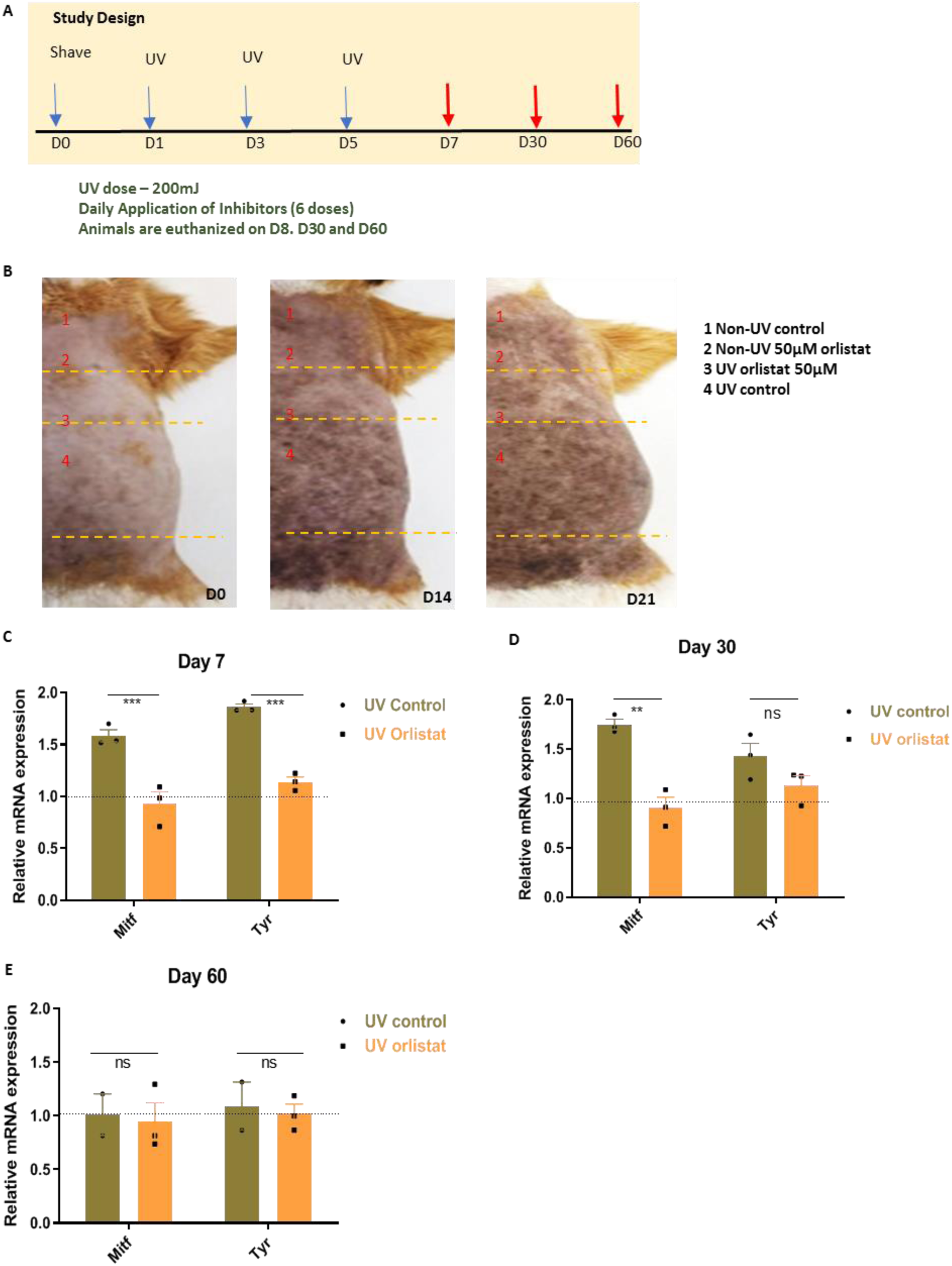
**Orlistat prevents induction of UV-mediated pigmentation in Guinea pigs** A. Experimental timeline showing UV-mediated pigmentation induction in guinea pigs. Three doses of UVA+B were given on day 1, 3 and 5 (shown by the blue arrow). Six doses of inhibitor were applied for the first six days. Animals were euthanized on D7, 30 and 60 (shown by the red arrow). B. Representative photographs showing phenotypic differences in guinea pig skin pigmentation on D14 and D21. Four segments were marked based on different treatments. These segments were compared across days. C. Bar graph depicting qRT-PCR based analysis of *Mitf* and *Tyr* expression from epidermal cells on day 7. Mean ± s.e.m. is plotted for three individual guinea pigs. One-way ANOVA is applied, F (3,8) = 36.64, p-value < 0.0001. Tukey’s test is performed for pairwise analysis. ***p-value < 0.005, **p-value < 0.01, *p-value < 0.05. D. Bar graph depicting qRT-PCR based analysis of *Mitf* and *Tyr* expression from epidermal cells on day 30. Mean ± s.e.m. is plotted for three individual guinea pigs. One-way ANOVA is applied, F (3,8) = 12.87, p-value = 0.002. Tukey’s test is performed for pairwise analysis. ***p-value < 0.005, **p-value < 0.01, *p-value < 0.05. E. Bar graph depicting qRT-PCR based analysis of *Mitf* and *Tyr* expression from epidermal cells on day 60. Mean ± s.e.m. is plotted for three individual guinea pigs. One-way ANOVA is applied, F (3,6) = 0.1190, p-value = 0.9457. Tukey’s test is performed for pairwise analysis. ns is not significant.

### Orlistat prevents induction of UV-mediated pigmentation in a Guinea pig model

Having established the important role of lipid metabolism in melanogenesis, we were interested to investigate whether the pharmacological targeting of fatty acid metabolism could show a therapeutic effect in an animal model. We established the guinea pig model to induce skin pigmentation, wherein 200 mJ of UVA and UVB dose were given for 3 alternate days (Fig 6A), as also reported previously ^41, 42^. Brown patches on guinea pig skin were analyzed in the study due to better contrast for hyper-pigmentation phenotype than the black patches. Since orlistat is an FDA approved drug, we examined whether topical application of orlistat could be a safe alternative for depigmentation. Orlistat was formulated with PEG8000 and applied topically at 50 µM for six days. As expected, we did not observe any toxicological effect of orlistat on the skin. We compared the phenotypic changes in the same animal by dividing the brown pigmented patch into 4 sections, which were subjected to different treatment conditions. The first section corresponds to the non-UV-exposed control region, while the second section is the non-UV-exposed orlistat treated region. The third section is the UV-exposed region with the application of the orlistat, and the fourth section is the UV-exposed control region. Phenotypic changes in the animals were captured on day 14 and day 21 by taking photographs (Fig 6B). Enhanced induction of pigment was clearly noted in section 4 due to UV exposure. Section 1 and 2, where the skin is not exposed to UV, show no changes in the pigmentation phenotype. In contrast, the orlistat-treated UV exposed region 3 showed reduced pigmentation as compared to the fourth quadrant. These phenotypic changes were confirmed at the molecular level on day 7, day 30, and day 60 when animals were euthanized. We observed that *Mitf* and *Tyr* expression increased upon UV treatment in the UV control region as compared to non-UV control, while UV orlistat treatment did not show a significant increase compared to the non-UV Orlistat set on day 7 and day 30 (Fig 6C, 6D). These molecular changes in various sections reversed after the recovery period of 60 days (Fig 6E). Together, this data clearly shows that the topical application of orlistat can be repurposed to decrease hyper-pigmentary responses and that modulators of metabolism may be a meaningful and safe way to regulate skin pigmentation.

## DISCUSSION

Seminal studies with infiltrating immune cells have underscored the importance of metabolism and metabolites as a guiding force and a critical determinant of the quality and quantity of immune responses ^43, 44^. However, how parenchymal cells like melanocytes rewire their metabolism to maintain cellular functions is largely unknown. Melanocytes in the human skin epidermis mostly consist of differentiated and non-dividing cells ^1^. These cells must rapidly and effectively respond to physiological cues like UV radiations and/or to the secreted factors like cytokines, growth factors and hormones and regain physiological homeostasis ^45, 46^. In this study, we have mapped the dynamics associated with transcriptional and metabolic networks that dictate the efficient melanogenesis response of melanocytes. To avoid confounding variables arising from different cell populations, we employed a functionally defined B16 cell pigmentation model that autonomously transits from basal depigmented to the pigmented state over a period of 6 days. This model recapitulates a series of coordinated processes encompassing signalling, transcriptional activation, melanosome biogenesis, melanin synthesis and return to a homeostatic state. Previous time-series analysis of this pigmentation model had resulted in the identification of interferon-γ signatures in dictating the depigmentation phase of melanogenesis ^26^. Our studies here delineate the pivotal role of fatty acid metabolism in melanocyte effector function and also establish the potential therapeutic efficacy of this target.

Integrated analysis of transcriptomic and metabolomics studies resolves melanocyte pigmentation function into three intricately synchronized phases corresponding to preparatory, melanogenic and recovery, defined by distinct transcriptional and metabolic signatures. During the preparatory phase, cells start to accumulate precursors and reducing equivalents by increasing the flux of glycolysis towards anabolic pathways like PPP and HBP. The shunting of glycolytic intermediates to the hexosamine biosynthesis module generates substrates for N-glycosylation that will facilitate extensive glycosylation of pigment-producing enzymes, TYR and DCT ^8, 47, 48^ and protein components of the melanosome like PMEL17 ^10^, which are heavily glycosylated in functional form. The flux through PPP remains high during the melanogenic phase that produces NADPH to facilitate the fatty acid biosynthesis crucial in the next phase. Increase phosphorylation of AKT during the preparatory phase of melanogenesis could sustain anabolic processes, increase glycolysis, and also enhances fatty acid synthesis ^49, 50^.

Induction of melanogenic phase activates fatty acid biosynthesis and accumulation of lipid droplets, which are then rapidly utilized for increased energy requirements of the cell. The onset of melanogenesis triggers gene expression of the central lipid mediator, SREBF1, and we observe activation of this TF maximally around the melanogenic phase. This activation is likely to trigger by various activators of pigmentation and we demonstrate α-MSH mediated cleavage and activation of SREBF1. Increased fatty acid synthesis results in the formation of TAGs. These lipid droplets are much smaller in size than those found in hepatocytes and are rapidly turned over during the continued melanogenic phase through β-oxidation ^51^. The accumulation of lipid droplets has been reported earlier within melanocytes in the skin of individuals exposed to therapeutic UV radiations ^52^. Several inhibitors targeting FASN, DGAT1, and lipase affect pigmentation phenotype, confirming that *de novo* fatty acid synthesis, storage, and lipolysis are integral in carrying out melanogenesis. Surprisingly, etomoxir-mediated inhibition of CPT1 does not show any effect on pigmentation (data not shown). While the reason for this data is somewhat puzzling, similar observations have been reported recently for regulatory and memory T cells ^53^.

Seahorse-based mitochondrial respiration analysis of these cells during pigmentation show that melanocytes have higher spare respiration capacity for utilizing fatty acids and thus have the potential to make use of this pathway during increased energy requirements. Mitochondrial reliance on fatty acids could result in increased accumulation of free radicals and ROS, which is manifested as the accumulation of defective mitochondria in the recovery phase. Our electron microscopy analysis demonstrates damaged mitochondria with distorted cristae surrounded by autophagic vacuoles in the later phase of melanogenesis. This corroborates with previous reports suggesting decreased mitochondrial respiration in hyper-pigmented cells ^21^. The switch to glycolysis and depletion of stored lipid droplets indicates an almost complete reliance on aerobic glycolysis for the cells’ energy needs. This is substantiated by an increased accumulation of lactate as a result of increased expression of PDK-1, which decreases pyruvate channelization to TCA. The NRF2 network may play a role in controlling the recovery phase, during which melanocytes restrict oxidative processes to attain homeostasis ^31^.

During pathological conditions, it is critical to selectively inhibit the pathways that dysregulate cellular functions such that treatment can restore homeostasis. Since the dynamic nature of metabolic programming amongst immune cells is linked with their plasticity and function, various inhibitors for metabolic pathways are being perused as a novel therapeutic approach to treat inflammation and autoimmunity^37, 43, 54^. However, one of the challenges is to generate selectivity during systemic administration for obtaining therapeutic benefits. In our study, by using inhibitors of fatty acid metabolism through a topical formulation, we are able to selectively target melanocyte function to treat hyper-pigmentary diseases. We showed that the FDA-approved drug, orlistat, can be a potential molecule to treat hyper-pigmentary responses. In conclusion, our study defines principles of cellular homeostasis during melanogenesis that reveals how melanocytes respond to systemic cues to elicit a physiological response by balancing energetic and cellular stability by sensing the environment. This knowledge can benefit in determining how cutaneous pigmentary diseases develop and as well as the means to treat them.

## MATERIALS AND METHODS

### B16 pigmentation model and primary melanocyte cultures

B16 melanoma cells were cultured in Dulbecco’s modified Eagle’s Media (DMEM-high glucose) supplemented with 10% fetal bovine serum (FBS). Cells were maintained at 5% CO_2_ levels at 37°C and grown till 60-80% confluency. For the pigmentation model, B16 cells were seeded at a low density of 100 cells/cm^2^ in DMEM-high glucose media and allowed to gradually pigment through different days of the model. Pigmentation was quantitated from pellet images using ImageJ software. Neonatal Human Epidermal Melanocytes (NHEM) were isolated from neonatal human foreskin samples and cultured in M254 media.

### UV-induced Guinea Pig pigmentation model

3-4-month-old guinea pigs were housed for the experiment. Animals were shaved with depilating cream to expose the skin a day before starting the UV dose. Three doses of 200 mJ of UVA+UVB was given on alternate days to half body of the animal while the remaining half was covered with aluminum foil, which serves as non-UV control skin ^41, 42^. Both the regions were then applied topically with an equal volume of 50 µM of Orlistat dissolved in 50% of polyethylene glycol 8000 (PEG8000). PEG8000 was used as vehicle control. In each experimental set, a total of six doses of drugs were given for the first six days. The first drug application was done after 1^st^ UV dose. Three animals were euthanized at each time point, day 7, 30 and 60. 1 X 1 cm^2^ of skin was collected from both control and treated region. Epidermis and dermis were separated by treating with 0.25% dispase solution in HBSS for 2 hours at 37°C. Transcriptional analysis was done for pigmentation genes from RNA isolated from the epidermis. UV exposed region was compared to the non-UV exposed region. All animal procedures were performed with an approved protocol from the institutional animal ethical committee at the National Institute of Immunology.

### RNA sequencing

RNA isolation was performed using the Macherey Nagel triprep kit according to the manufacturer’s instructions. Briefly, B16 cells were cultured at low density on consecutive days such that day 6 and day 3-time points would coincide on the same day. For RNA isolation, an equal number of cells (5X10^5^) were counted for each time point and stored in the lysis buffer. To obtain high-quality RNA, purification was performed using triprep column. The quality of extracted RNA was assessed by visualizing bands on 1.5% agarose gel and monitoring 260/280 ratio. All the RNA samples were frozen together at -80°C. RNA was precipitated with ethanol to preserve the quality during transportation. 1µg RNA sample was sent for two replicates of each day for RNA sequencing. RNA samples were outsourced to the company, Bencos Research Solution Pvt Ltd, for RNA sequencing. RNA with RIN > 8.5 was proceeded for library preparation using TruSeq RNA Sample Prep Kit v2. Sequencing was performed using Illumina NovaSeq 6000 platform. QC check was done using FastQC. Pre- processing to remove trim bases and adaptor was done using Cutadapt. Read alignment was done using HISAT2. GRCm38 was used as the reference genome. Read quantification was performed with HTSeq and normalized counts and differential regulation were obtained using the DESeq2 package in R. Time course analysis was done using the LRT test and genes with adjusted p-value < 0.001 were taken as significantly differentially expressed from day3 to day6. Raw counts were transformed using the vst function and DEG’s were clustered and plotted using the pheatmap package. Genes enriched on the respective days were identified and KEGG pathway enrichment was done using the clusterProfiler package. Selected pathways were plotted using ggplot2 in R Studio. DEG’s were analysed for the target (TG) enrichment of transcription factors (TF) using the TRUSST database on metascape (metascape.org). Top 7 transcription factors from each day (except Trp53 as it was enriched on all the days) were plotted as a TF-TG network using cytoscape. The colour gradient represents the scaled (row-wise) expression values from the RNA Seq data.

### cDNA synthesis and real-time PCR

RNA was reverse transcribed using the Superscript III cDNA synthesis kit (Life Technologies) according to the manufacturer’s protocol. Gene expression analysis by quantitative real-time PCR was performed on a Roche Light Cycler 480 II real-time cycler using the SYBR GREEN FAST qPCR Master Mix (Thermo) to evaluate transcriptional regulations. Most of the primers were designed using Primer3 and checked by the NCBI Primer blast tool. Gene-specific primers were obtained from Sigma Aldrich. Either *Hgprt or Gapdh* was used as the normalizing control and quantification was done by the comparative Ct method.

**Table 1.**
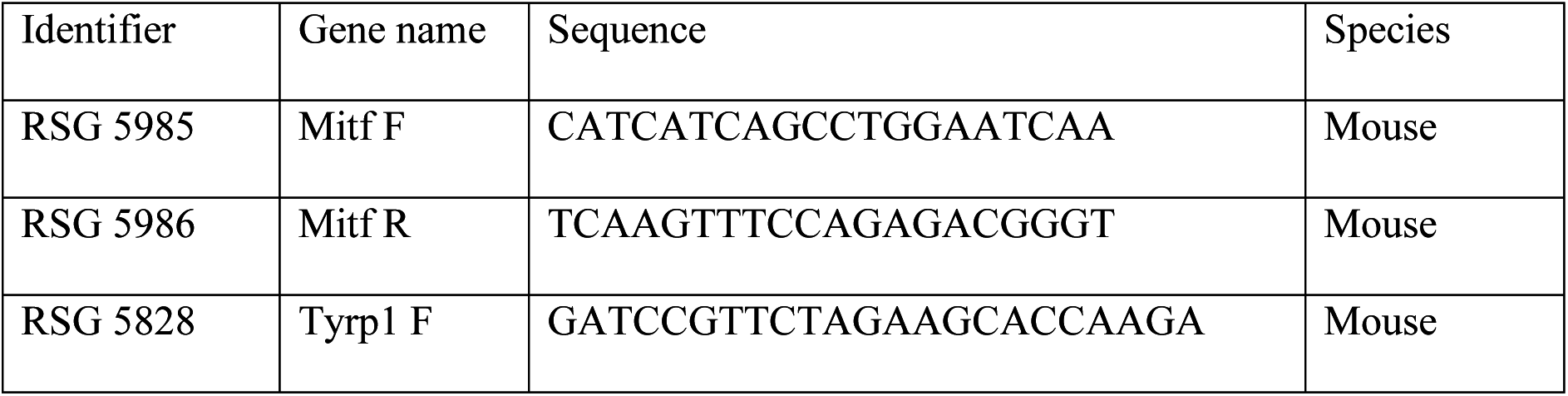
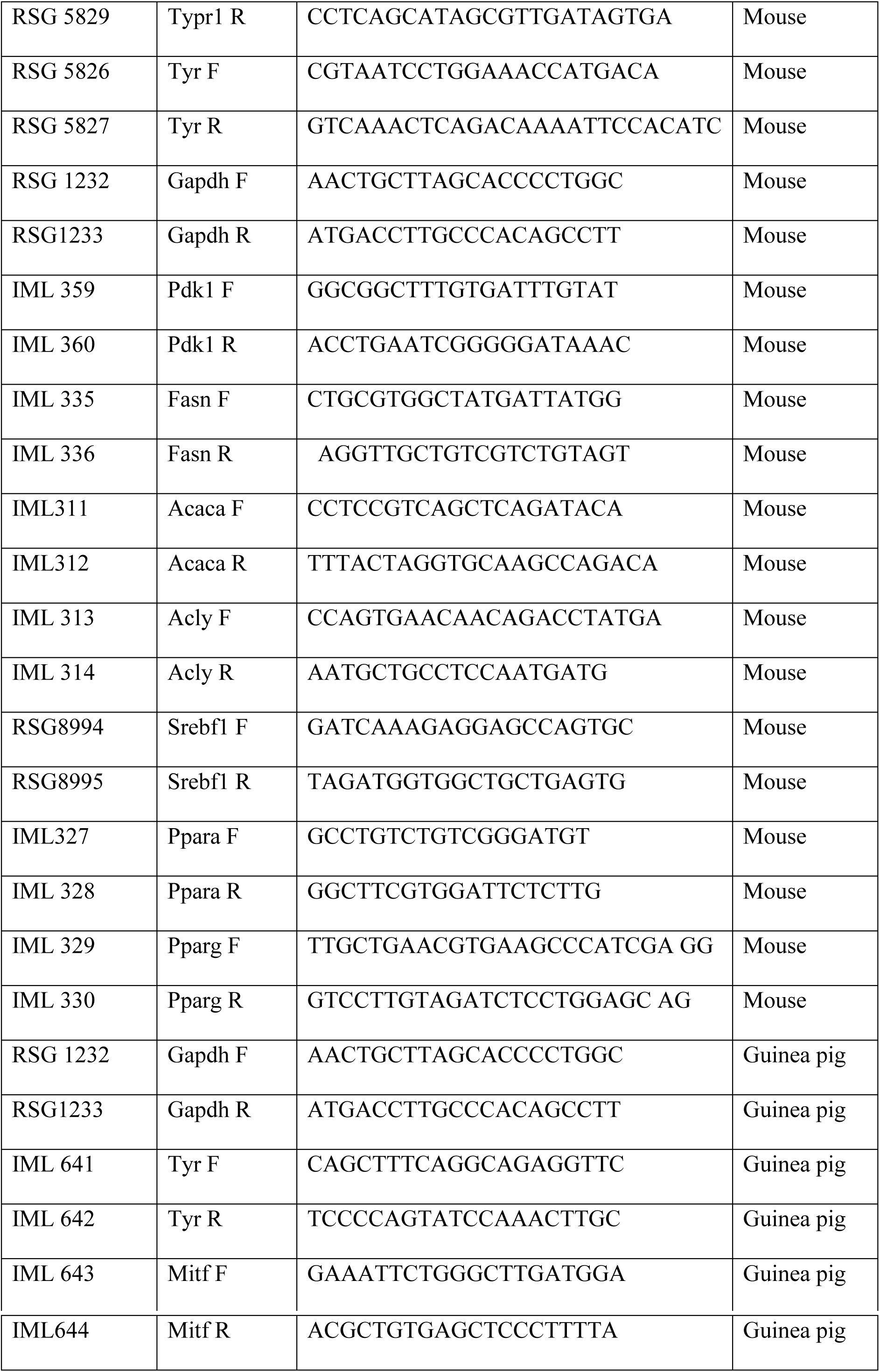
List of qRT-PCR primers

### Cell lysate preparation and western blotting

B16 melanoma cells were washed with ice-cold Phosphate buffer saline (PBS) and lysed with NP40 lysis buffer supplemented with protease inhibitor cocktail. Cells were incubated with NP40 for 30 minutes on ice. The soluble fractions of cell lysates were isolated by centrifugation at 13,000 rpm in a refrigerated microcentrifuge for 30 minutes. The protein concentration in the soluble fraction was quantified using the bicinchoninic acid (BCA) protein estimation kit. Known concentrations of bovine serum albumin (BSA) was used to plot the standard curve. 30-50 μg of the protein was boiled in SDS dye and separated on 10% SDS PAGE gel. Tyrosinase antibody is synthesized from Genescript. DCT (ab74073), PMEL17 (ab137078), MITF (ab12039), FASN (ab22759) and mCHERRY (ab167453) antibodies are procured from Abcam, CDK2 (MA1-81135) is obtained from Thermo Scientific while Srebf1 (04-469) is ordered from Millipore. HRP-conjugated Actin (ab8227) and Tubulin (ab6046) are used as a loading control. Horseradish peroxidase-conjugate Anti-Mouse (NA931) and Anti-Rabbit (NA934) antibodies are obtained from GE healthcare. For Western blot standard enhanced chemiluminescence reagents (WBLUF0100) were used from Millipore. ImageJ software was used for quantification.

### Glucose measurement

Glucose concentration in media was measured on COBAS INTEGRA 400 using a glucose detection kit. It is based on the principle of enzymatic assay where glucose is first converted to glucose-6-phosphate by hexokinase, followed by oxidation of glucose-6-phosphate by G6PD, which is coupled to the reduction of NAD to NADH. NADH produced in the reaction reduces colorless probe to a colored product with strong absorbance at 450nm. The amount of glucose is equivalent to NADH produced. A standard curve was generated using glucose standards ranging in between 0-10 g/L concentration. The instrument is highly accurate up to the range of 6 g/L. 200 µl residual media was collected on each day from day 3-day 6 in specialized tubes and placed in the COBAS instrument. Reagents were mixed automatically and readings were recorded. Glucose concentration in the media on different days was calculated from the standard curve.

### Immunofluorescence measurements

Lipid droplets were measured using BODIPY dye. B16 cells were grown on 1cm^2^ coverslip in low density (1000 cells per coverslip) for different days. Cells were fixed by adding 4% methanol-free formaldehyde on the coverslip and incubated at room temperature for 20 minutes. 3X PBS washes were given to remove 4% methanol-free formaldehyde. Cells were stored in PBS at 4°C till the last time point. Fixed cells were incubated with 10 µM BODIPY dye in DPBS for 1 hour at room temperature. 3X PBS washes were done to remove excess stain. Coverslip was mounted with Gold antifade DAPI solution. BODIPY staining in primary cells was done without fixation. In primary melanocytes, fixation was leading to coalesce of lipid droplets due to their bigger size. Live primary melanocytes were incubated with 10µM BODIPY dye solution in HBSS for 40 minutes at 37°C. Imaging was done on the Leica SP8 confocal microscope. Three biological replicates were imaged for each experiment, with 50 cells per set in the B16 model.

B16 cells overexpressing pmCherry-Srebf1-eGFP (traffic light construct) were seeded in live imaging chambers (Two chambered live imaging chambers from Nunc). 3.5 μM Insulin or 10 μM α-MSH was added just before setting up the live imaging module in the Leica SP8 Confocal microscope. Transfected cells were selected at random using the “Mark & Find” feature and 40-50 cells were imaged for a duration of 4-6 hours per replicate. The experiment was performed in triplicates.

### Polar metabolite extraction for Mass spectrometry and NMR

For liquid chromatography-coupled tandem mass spectrometric analysis, polar metabolite extraction was performed using 80% Methanol-Water solvent. Cells were grown in multiple flasks to obtain 10^6^ cells per replicate. 3-4 replicates were prepared for each time point. B16 cells were washed with cold PBS and trypsinized with 0.1% trypsin. Cells were harvested in defined trypsin inhibitor, and one million cells were counted for each sample and proceeded for metabolite extraction. Again, cells were washed with PBS and pelleted down at 500 g for 5min at 4°C. 500 µL of chilled 80% methanol was added to the cell pellet. For extraction, samples were mixed well by vortexing and incubated on dry ice for 10 minutes. Penicillin was added in 80% methanol as an internal control during extraction. Samples were spun at 13000 rcf for 30 min at 4°C. The supernatant was collected in a chilled 1.5 ml micro-centrifuge tube and samples were dried in a SpeedVac vacuum concentrator. Samples were further lyophilized and stored at -80°C till mass spectrometric run. The samples were run on a Waters Xevo-TQS tandem mass spectrometer coupled to Acquity UPLC. The analysis was performed using MassLynx software, followed by analysis with MetaboAnalyst 4.0 for pathway enrichment. Fold changes were plotted for each metabolite.

Pulse-chase labeling of glycolysis and TCA metabolites was performed using NMR. 10 mM [U-^13^C]-Glucose was fed in glucose-free RPMI media for 4 hours at each time point. Due to the low sensitivity of NMR, extraction was done from 20 million cells per sample. 200 µM [U- ^13^C]-palmitate was added for 24 hours in glucose-free RPMI at each time point. Metabolite extraction was done with the same protocol as mention above. Samples were dissolved in 160 µL of deuterated water (D_2_O) and transferred in 3mm NMR tubes. All NMR measurements were performed on a 500 MHz Bruker Avance III spectrometer equipped with 5 mm TCI cryo- probe. Topspin 3.6 pl7 (Bruker) was used for data acquisition, Fourier transformation and processing of data. Two-dimensional [^13^C,^1^H] heteronuclear single quantum coherence [HSQC] experiments were measured at 310 K. The 2D [^13^C,^1^H] HSQC, spectra were measured with a spectral width of 7002.8 Hz along the ^1^H dimension, and 22639.57 Hz along the ^13^C dimension. A total of 16 dummy scans and 96 scans with a relaxation delay of 1.5 sec was used for a total acquisition time of 146 ms (_t2max_) along ^1^H dimension and 2.8 ms (t_1max_) along ^13^C dimension. Processing was performed using 90° shifted sine-square bell window function for both dimensions. Peak correlation and peak intensity calculation were performed using Computer Aided Resonance Assignment (CARA) software ^55^. Chemical shift values were assigned to specific metabolites using the Biological Magnetic Resonance Bank (BMRB) (http://www.bmrb.wisc.edu) and Human Metabolome Database (HMDB) (http://www.hmdb.ca). A chemical shift error-tolerance of 0.05 ppm and 0.5 ppm was used for ^1^H and ^13^C chemical shifts, respectively. Fold change as compared with day 0 was plotted for each metabolite.

### Lactate measurement

Lactate was measured in cell lysate with the Sigma Lactate assay kit (MAK064) using the manufacturer’s protocol. Briefly, an equal number of cells (10^6^) were harvested on day 5 and day 6 and the cellular lysate was prepared using the given lysis buffer. Lactate standards were prepared in a range of 0-10 nmole in 50 µL of Assay buffer. An equal volume of a master mix containing lactate assay buffer, lactate enzyme and lactate probe were added to standards and samples. The plate was incubated for 30 minutes at room temperature and colorimetric measurement was recorded at 570nm.

### siRNA transfection

siRNA transfections were performed in T75 flasks on day 3 of the pigmentation model. 100nM of siRNA was added per flask with a 1: 3 times V: V ratio of Dharmafect transfection reagent (SiRNA catalogue provided in supplementary information). The transfection was done in opti-MEM media for 6 hours. Post transfection, the opti-MEM media was removed and cells were washed with 1X DPBS and then the day 3 culture media was added back to the cells.

### Melanin estimation

Melanin estimation was performed as described earlier ^56^. Briefly, cells were lysed in 1 N NaOH by heating at 80°C for 2 h, and then, absorbance was measured at 405nm. Finally, the melanin content was estimated by interpolating the sample readings on the melanin standard curve obtained with synthetic melanin.

### Cloning

To clone *Srebf1*, we amplified the common splice variant, *Srebf1c*, using PCR. To get a complete 3.4 kb fragment, we amplified the first 1.6 kb separately and the last 1.7 kb sequence separately. The complete sequence was amplified with overlapping PCR by using two amplicons specific to the first and second half sequence of *Srebf1c*. The amplicon was phosphorylated with T4 polynucleotide kinase. The plasmid pBluescript (pBS-SK+) was digested with EcoRV and end were dephosphorylated with calf intestinal phosphatase (CIP). Amplified full-length sequence of *Srebf1c* was cloned in pBS-SK+ vector in EcoRV restriction digestion site. From Srebf1-pBS plasmid, *Srebf1c* fragment was removed by doing partial restriction digestion with XhoI and EcoRI enzymes. Complete digestion gives two fragments, from 1-493 bp and 494 to 3410 bp, while partial digestion gives three bands corresponding full length 1-3410 bp and 1-493 bp and 494 to 3410 bp. We extracted the full-length fragment from agarose gel corresponding to 3335 bp and sub-cloned it in the mCherry-C1 vector. In our lab, eGPF was already cloned in the mCherry-C1 vector and stored in lab repository as clone number pAK4.0. With XhoI and EcoRI restriction enzymes, we sub-cloned *Srebf1c* in mCherry-eGFP vector.

**Table 2.**
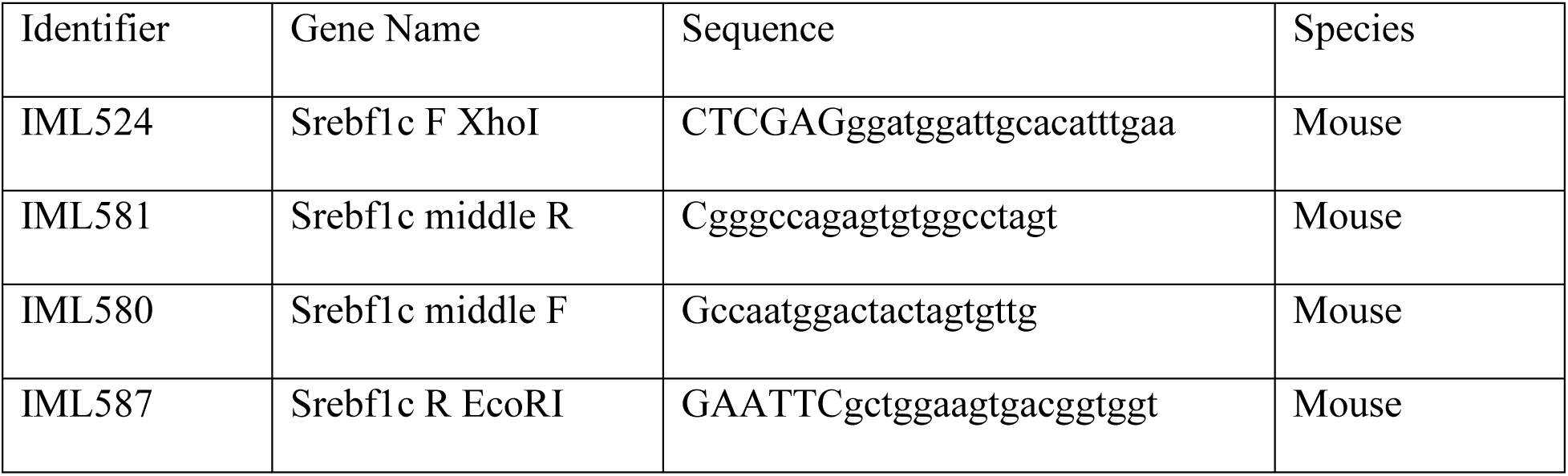
List of cloning primers

### Transmission Electron Microscopy

Cells were fixed in fixative containing 2.5% glutaraldehyde and 4% paraformaldehyde in 0.1 M sodium cacodylate buffer (pH 7.2) at room temperature for 4 hours and then rinsed in PBS. Fixed cells were embedded in 2% agar and post-fixed in 1% osmium tetraoxide for 1 hour. Samples were dehydrated in graded series of ethanol (50–100%) and then subjected to propylene-oxide for 30 minutes, and infiltrated with increasing proportions of propylene-oxide: Epon (2:1, 1:1 and 1:2) for 2 hours and embedded in Epon resin and polymerized for 72 hours at 60°C. Ultrathin sections (63 nm) were cut on Leica Ultra-microtome, placed on copper grids and stained with uranyl acetate and lead citrate, and examined on a 200 KV Tecnai G2 Twin transmission electron microscope (FEI make).

### Statistical analysis

All statistical analysis was performed using GraphPad Prism 8 software. Unpaired student’s t-test, one-way ANOVA or two-way ANOVA was applied to compare the significant difference between the means of multiple groups. Tukey’s test was applied to compare the pair-wise sample. Dunnett’s test was applied to compare with the control group. * represents p-value ≤ 0.05, ** represents p-value ≤ 0.01 and *** represents p-value ≤ 0.001. The p-values obtained during analysis are indicated in the figure legends.

## Data Availability

The accession number for the RNA sequencing data reported in this paper is GSE164375. Further, the authors declare that the other data supporting the findings of this study are available within the paper and its Supplementary Information files.

## Acknowledgement

R.S.G. acknowledges support from J.C. Bose fellowship and Department of Biotechnology (DBT) for providing funds to National Institute of Immunology. R.S.G., T.N.V. & C.G. acknowledge the support of CSIR project TOUCH (BSC0302). F.S. acknowledges UGC JRF/SRF fellowship. D.M. and K.S.A. were CSIR JRF/SRF Fellows. M.K. was supported by funds from SERB project. P.K., P.D. & A.A. are recipients of the CSIR JRF/SRF fellow. We acknowledge the infrastructure supported by CSIR-IGIB for imaging facility and Mr. Manish for help with imaging. We thank Department of Biotechnology, Government of India and ICGEB, New Delhi core research fund for providing financial support for the high field NMR spectrometer at ICGEB, New Delhi.

## Authors contribution

R.S.G., T.N.V. and F.S. conceived the project, designed the experiments, analysed data and wrote the paper. D.M. was involved in initiating the study. F.S., R.B. and M.K. performed experiments and participated in the discussion of the results. F.S. and M.K. assisted with animal experiments. C.G., P.D. and A.A. assisted in data analysis. K.S.A. and P.K.S. contributed to mass spectrometry data generation. N.S.B. and P.K. contributed to NMR experiments. A.S. performed transmission electron microscopy.

## References

1. Costin, G.-E. & Hearing, V. J. Human skin pigmentation: melanocytes modulate skin color in response to stress. FASEB J. 21, 976–994 (2007).

2. Yosipovitch, G., DeVore, A. & Dawn, A. Obesity and the skin: skin physiology and skin manifestations of obesity. J. Am. Acad. Dermatol. 56, 901–916 (2007).

3. Duff, M., Demidova, O., Blackburn, S. & Shubrook, J. Cutaneous manifestations of diabetes mellitus. Clin. Diabetes, 33, 40–48. (2015).

4. Cichorek, M. & Wachulska, M. Skin melanocytes: biology and development. Postep. Dermatol Alergol 30, 30–41 (2013).

5. Ando, H. et al. Melanosomes are transferred from melanocytes to keratinocytes through the processes of packaging, release, uptake, and dispersion. J. Investig. Dermatology, 132, 1222–1229 (2012).

6. Gibbs, S. et al. Melanosome capping of keratinocytes in pigmented reconstructed epidermis–effect of ultraviolet radiation and 3-isobutyl-1-methyl-xanthine on melanogenesis. Pigment cell Res. 13, 458–466 (2000).

7. Goding, C. R. & Arnheiter, H. MITF—the first 25 years. Genes Dev. 33, 983–1007. (2019).

8. Tsukamoto, K., Jackson, I. J., Urabe, K., Montague, P. M. & Hearing, V. J. A second tyrosinase-related protein, TRP-2, is a melanogenic enzyme termed DOPAchrome tautomerase. journal,. EMBO 11, 519–526. (1992).

9. Jimbow, K. et al. Assembly, target-signaling and intracellular transport of tyrosinase gene family proteins in the initial stage of melanosome biogenesis. Pigment Cell Res. 13, 222–229. (2000).

10. Berson, J. F., Harper, D. C., Tenza, D., Raposo, G. & Marks, M. S. Pmel17 initiates premelanosome morphogenesis within multivesicular bodies. Mol. Biol. cell, 12, 3451–3464. (2001).

11. McGill, G. G. et al. Bcl2 regulation by the melanocyte master regulator Mitf modulates lineage survival and melanoma cell viability. Cell, 109, 707–718. (2002).

12. Du, J. et al. Critical role of CDK2 for melanoma growth linked to its melanocyte-specific transcriptional regulation by MITF. Cancer cell, 6, 565–576. (2004).

13. Robbins, L. S. et al. Pigmentation phenotypes of variant extension locus alleles result from point mutations that alter MSH receptor function. Cell, 72, 827–834 (1993).

14. Bertolotto, C. et al. Microphthalmia gene product as a signal transducer in cAMP-induced differentiation of melanocytes. J. cell Biol. 142, 827–835. (1998).

15. Chakraborty, A. K. et al. Production and release of proopiomelanocortin (POMC) derived peptides by human melanocytes and keratinocytes in culture: regulation by ultraviolet B. Biochim. Biophys. Acta (BBA)-Molecular Cell Res. 1313, 130–138. (1996).

16. Rousseau, K. et al. Proopiomelanocortin (POMC), the ACTH/melanocortin precursor, is secreted by human epidermal keratinocytes and melanocytes and stimulates melanogenesis. The FASEB Journal, 21, 1844–1856. (2007).

17. Bondurand, N. et al. Interaction among SOX10, PAX3 and MITF, three genes altered in Waardenburg syndrome. Hum. Mol. Genet. 9, 1907–1917. (2000).

18. Furumura, M. et al. Involvement of ITF2 in the transcriptional regulation of melanogenic genes. J. Biol. Chem. 276, 28147–28154 (2001).

19. Ferguson, J., Smith, M., Zudaire, I., Wellbrock, C. & Arozarena, I. Glucose availability controls ATF4-mediated MITF suppression to drive melanoma cell growth. Oncotarget, 8, 32946. (2017).

20. Rachmin, I., Ostrowski, S. M., Weng, Q. Y. & Fisher, D. E. Topical treatment strategies to manipulate human skin pigmentation. Adv Drug Deliv Rev 01, 65–71 (2020).

21. Meira, W. V., Heinrich, T. A., Cadena, S. M. S. C. & Martinez, G. R. Melanogenesis inhibits respiration in B16-F10 melanoma cells whereas enhances mitochondrial cell content. Exp. Cell Res. 350, 62–72 (2017).

22. Daniele, T. et al. Mitochondria and melanosomes establish physical contacts modulated by Mfn2 and involved in organelle biogenesis. Curr. Biol. 24, 393–403. (2014).

23. Kim, E. S. et al. Mitochondrial dynamics regulate melanogenesis through proteasomal degradation of MITF via ROS-ERK activation. Pigment Cell Melanoma Res. 27, 1051–1062 (2014).

24. Jung, D. W. et al. Identification of the F1F0 mitochondrial ATPase as a target for modulating skin pigmentation by screening a tagged triazine library in zebrafish. Mol. Biosyst. 1, 85–92. (2005).

25. Seo, S. H. et al. Metabolomics reveals the alteration of metabolic pathway by alpha-melanocyte-stimulating hormone in B16F10 melanoma cells. Molecules 25, (2020).

26. Natarajan, V. T. et al. IFN-γ signaling maintains skin pigmentation homeostasis through regulation of melanosome maturation. Proc. Natl. Acad. Sci. 111, 2301–2306. (2014).

27. Malcov-Brog, H. et al. UV-Protection Timer Controls Linkage between Stress and Pigmentation Skin Protection Systems. Mol. Cell 72, 444–456.e7 (2018).

28. Shin, S.Y., Choi, J.H., Jung, E., Gil, H.N., Lim, Y. and Lee, Y. H. The EGR1-STAT3 Transcription Factor Axis Regulates α-Melanocyte-Stimulating Hormone-Induced Tyrosinase Gene Transcription in Melanocytes. The Journal of investigative dermatology,. J. Invest. Dermatol. 139, 1616–1619 (2019).

29. Wang, H., Mannava, S. Grachtchouk, V. Zhuang, D., Soengas, M. S., Gudkov, A. V. & Prochownik, E.V. Nikiforov, M. A. c-Myc depletion inhibits proliferation of human tumor cells at various stages of the cell cycle. Oncogene 27, 1905–1913 (2008).

30. Inoue, J. et al. Proteolytic activation of SREBPs during adipocyte differentiation. Biochem. Biophys. Res. Commun. 283, 1157–1161. (2001).

31. Natarajan, V. T. et al. Transcriptional upregulation of Nrf2-dependent phase II detoxification genes in the involved epidermis of vitiligo vulgaris. J. Invest. Dermatol. 13-, 2781–2789 (2010).

32. Ando, H. et al. Fatty acids regulate pigmentation via proteasomal degradation of tyrosinase: a new aspect of ubiquitin-proteasome function. J Biol Chem. 9, 15427–33 (2004).

33. Semba, H. et al. HIF-1α-PDK1 axis-induced active glycolysis plays an essential role in macrophage migratory capacity. Nat. Commun. 7, 11635. (2016).

34. Kim, J. W., Tchernyshyov, I., Semenza, G. L. & Dang, C. V. HIF-1-mediated expression of pyruvate dehydrogenase kinase: a metabolic switch required for cellular adaptation to hypoxia. Cell Metab. 3, 177–185. (2006).

35. Filipp, F. V et al. Glutamine-fueled mitochondrial metabolism isdecoupled from glycolysis in melanoma. Pigment Cell Melanoma Res. 25, 732–739 (2012).

36. Aloia, A. et al. A fatty acid oxidation-dependent metabolic shift regulates the adaptation of BRAF-mutated melanoma to MAPK inhibitors. Clin. Cancer Res. 25, 6852–6867 (2019).

37. Pan, Y. et al. Survival of tissue-resident memory T cells requires exogenous lipid uptake and metabolism. Nature 543, 252–256 (2017).

38. Kalucka, J. et al. Quiescent Endothelial Cells Upregulate Fatty Acid β-Oxidation for Vasculoprotection via Redox Homeostasis. Cell Metab. 28, 881–894.e13 (2018).

39. Buck, M. D. D. et al. Mitochondrial Dynamics Controls T Cell Fate through Metabolic Programming. Cell 166, 63–76 (2016).

40. Dif, N. et al. Insulin activates human sterol-regulatory-element-binding protein-1c (SREBP-1c) promoter through SRE motifs. *Biochem*. Journal, 400, 179–188. (2006).

41. Yoshida, M., Hirotsu, S., Nakahara, M., Uchiwa, H. & Tomita, Y. Histamine is involved in ultraviolet B-induced pigmentation of guinea pig skin. J. Investig. dermatology, 118, 255–260. (2002).

42. Allan, A. E., Archambault, M., Messana, E. & Gilchrest, B. A. Topically applied diacylglycerols increase pigmentation in guinea pig skin. J. Invest. Dermatol. 105, 687–692 (1995).

43. Berod, L. et al. De novo fatty acid synthesis controls the fate between regulatory T and T helper 17 cells. Nat. Med. 20, (2014).

44. Everts, B. J. A. K. H. S. C.-C. S. A. L. V. I. Y. L. E. C. K. S. K. M. A. J. Network integration of parallel metabolic and transcriptional data reveals metabolic modules that regulate macrophage polarization. Immunity 42, 419–430 (2015).

45. Natarajan, V. T., Ganju, P., Ramkumar, A., Grover, R. & Gokhale, R. S. Multifaceted pathways protect human skin from UV radiation. Nature Chemical Biology vol. 10 542–551 (2014).

46. Imokawa, G., Yada, Y., Kimura, M. & Morisaki, N. Granulocyte/macrophage colony-stimulating factor is an intrinsic keratinocyte-derived growth factor for human melanocytes in UVA-induced melanosis. Biochem. J. 313, 625–631 (1996).

47. Negroiu, G., Branza-Nichita, N., Petrescu, A. J., Dwek, R. A. & Petrescu, S. M. Protein specific N-glycosylation of tyrosinase and tyrosinase-related protein-1 in B16 mouse melanoma cells. . *Biochem*. journal, 344, 659–665 (1999).

48. Mikami, M. et al. ‘Glycosylation of tyrosinase is a determinant of melanin production in cultured melanoma cells’. Mol. Med. 8, 818–822 (2013).

49. Yecies, J. L. et al. Akt stimulates hepatic SREBP1c and lipogenesis through parallel mTORC1-dependent and independent pathways. Cell Metab. 14, 21–32 (2011).

50. Cheng, J. et al. TRIM21 and PHLDA3 negatively regulate the crosstalk between the PI3K/AKT pathway and PPP metabolism. Nat. Commun. 11, 1–16 (2020).

51. Suzuki, M., Shinohara, Y., Ohsaki, Y. & Fujimoto, T. Lipid droplets: Size matters. J. Electron Microsc. (Tokyo). 60, 101–116 (2011).

52. Schuler, G., Hönigsmann, H., Jaschke, E. & Wolff, K. Selective accumulation of lipid within melanocytes during photochemotherapy (PUVA) of psoriasis. Br. J. Dermatology, 107, 173–182. (1982).

53. Raud, B. et al. Etomoxir Actions on Regulatory and Memory T Cells Are Independent of Cpt1a-Mediated Fatty Acid Oxidation. Cell Metab. 28, 504–515.e7 (2018).

54. O’Sullivan, D. et al. Memory CD8+ T Cells Use Cell-Intrinsic Lipolysis to Support the Metabolic Programming Necessary for Development. Immunity 41, 75–88 (2014).

55. Keller, R. & Wuthrich, K. Computer-aided resonance assignment (CARA). Verl Goldau Cantina Switz. (2004).

56. Kageyama, A. et al. Down-regulation of melanogenesis by phospholipase D2 through ubiquitin proteasome-mediated degradation of tyrosinase. J. Biol. Chem. 279, 27774–27780. (2004).

